# Phage-bacteria dynamics during the first years of life revealed by trans-kingdom marker gene analysis

**DOI:** 10.1101/2023.09.28.559994

**Authors:** Michael Tisza, Richard Lloyd, Kristi Hoffman, Daniel Smith, Marian Rewers, the TEDDY Study Group, Sara Javornik Cregeen, Joseph F. Petrosino

**Affiliations:** The Alkek Center for Metagenomics and Microbiome Research, Department of Molecular Virology and Microbiology, Baylor College of Medicine, Houston, TX 77030, USA; Department of Molecular Virology and Microbiology, Baylor College of Medicine, Houston, TX, USA; Barbara Davis Center for Childhood Diabetes, University of Colorado, Aurora, CO, USA

## Abstract

Humans are colonized with commensal bacteria soon after birth, and, while this colonization is affected by lifestyle and other factors, bacterial colonization proceeds through well-studied phases. However, less is known about phage communities in early human development due to small study sizes, inability to leverage large databases, and lack of appropriate bioinformatics tools. In this study, whole genome shotgun sequencing data from the TEDDY study, composed of 12,262 longitudinal samples from 887 children in 4 countries, is reanalyzed to assess phage and bacterial dynamics simultaneously. Reads from these samples were mapped to marker genes from both bacteria and a new database of tens of thousands of phage taxa from human microbiomes. We uncover that each child is colonized by hundreds of different phages during the early years, and phages are more transitory than bacteria. Participants’ samples continually harbor new phage species over time whereas the diversification of bacterial species begins to saturate. Phage data improves the ability for machine learning models to discriminate samples by country. Finally, while phage populations were individual-specific, striking patterns arose from the larger dataset, showing clear trends of ecological succession amongst phages, which correlated well with putative host bacteria. Improved understanding of phage-bacterial relationships may reveal new means by which to shape and modulate the microbiome and its constituents to improve health and reduce disease, particularly in vulnerable populations where antibiotic use and/or other more drastic measures may not be advised.

## Introduction

From birth, the guts of all healthy humans play host to commensal bacteria^1,2^. These bacteria extensively metabolize our dietary inputs^3,4^, drive normal development of the immune system^5^, and, to our detriment, include some microbes that act as deadly opportunistic pathogens^6^.

Commensal bacteria are also hosts to (bacterio)phages, which are viruses that infect bacterial cells. These come in two main types: “virulent phages” which quickly replicate and lyse their hosts and “temperate phages” which integrate into the host cells as prophage, typically for many generations, before lysing the bacteria upon some cue or stressor^7^. Because phages exert pressure on host cell populations, and many phage genomes encode virulence factors and toxins^8^, a role for specific phages or communities of phages in human health seems plausible.

Previous studies have begun to explore how communities of bacteria or phages develop in infants and young children. Bacterial studies have shown that nearly all infants are initially contacted and/or colonized by bacteria prenatally or during delivery^2,9^. In the first days of life, bifidobacteria and other bacteria capable of efficiently metabolizing breast milk and formula dominate^10,11^. Then, human developmental stages determine bacterial community structure with a handful of taxonomic compositions frequently observed at different ages^12^. Studies of phages in infants are more limited but have shown that the first recoverable phages are typically prophages induced from early-colonizing bacteria^13^. Next, phage diversity and abundance increase and more virulent phages can be observed^14^. In infants and adults, phage populations have been shown to be more individual-specific than bacteria, making further trends and patterns difficult to uncover^15,16^. Therefore, answering how phage population dynamics are related to bacterial population dynamics, and whether bacterial and phage developmental phases are similarly deterministic, are among the questions that have eluded the field thus far.

In order to answer these questions, we built an extensive phage genome catalog from several recent gut phage meta-studies^17–20^ from which we extracted unique marker genes with essential phage functions and added them to the MetaPhlAn4 bacterial marker gene database^21^. We used this combined phage-bacteria marker gene database for the simultaneous profiling of phage and bacteria in 12,262 longitudinal stool samples from 887 participants in the TEDDY study^22,23^.

Longitudinal analysis showed that phage communities change more quickly than bacterial communities, with most phages persisting in a participant for a shorter duration. Subsequently, each participant hosted a more diverse repertoire of phages than bacteria during early development. Despite this, patterns of ecological succession were observed in the data, with different phages peaking in abundance at different host ages, largely mirroring the abundance of putative host bacteria. Adding phage taxonomic profiles improved the ability to discriminate samples geographically over bacteria taxonomic profiles alone. Furthermore, modest differences in phage and bacteria communities were observed in participants diagnosed with type 1 diabetes.

## Results

### Simultaneous profiling of viruses and bacteria in whole genome shotgun sequencing data via the Marker-MAGu pipeline

To enable accurate profiling of phages in whole genome shotgun (WGS) sequence datasets from stool samples, a more comprehensive database of gut phages (and a small number of eukaryotic viruses) was compiled from publicly available resources^17–20^. This database, the Trove of Gut Virus Genomes (see Materials & Methods), consists of genomes from 110,296 viral Species-level Genome Bins (SGBs)^24^ derived from human gut metagenome studies. Genomes from 42.6% of the viral SGBs are predicted to be 90 - 100% complete, and the database contains numerous phages infecting all major taxa of gut bacteria (Fig S1).

To facilitate comparable detection of viral and cellular genomes in WGS data, we developed a marker gene approach. This approach utilizes the concept that some genes are more taxonomically informative than others and, therefore, requires representing genomes by a subset of genes that are species-specific and invariable. For each viral genome in Trove of Gut Virus Genomes, essential genes (i.e., those involved in virion structure, genome packaging, and genome replication) were annotated as potential markers. After dereplication, 416,428 unique viral marker genes were detected. The 49,111 virus genomes with four or more unique marker genes were used for taxonomic profiling. We developed a bioinformatics tool, Marker-MAGu, leveraging marker genes and taxonomic identities from MetaPhlAn4 and Trove of Gut Virus Genomes to generate trans-kingdom taxonomic profiles for gut metagenomes (Fig 1A-B) (https://github.com/cmmr/Marker-MAGu).

**Figure 1.**
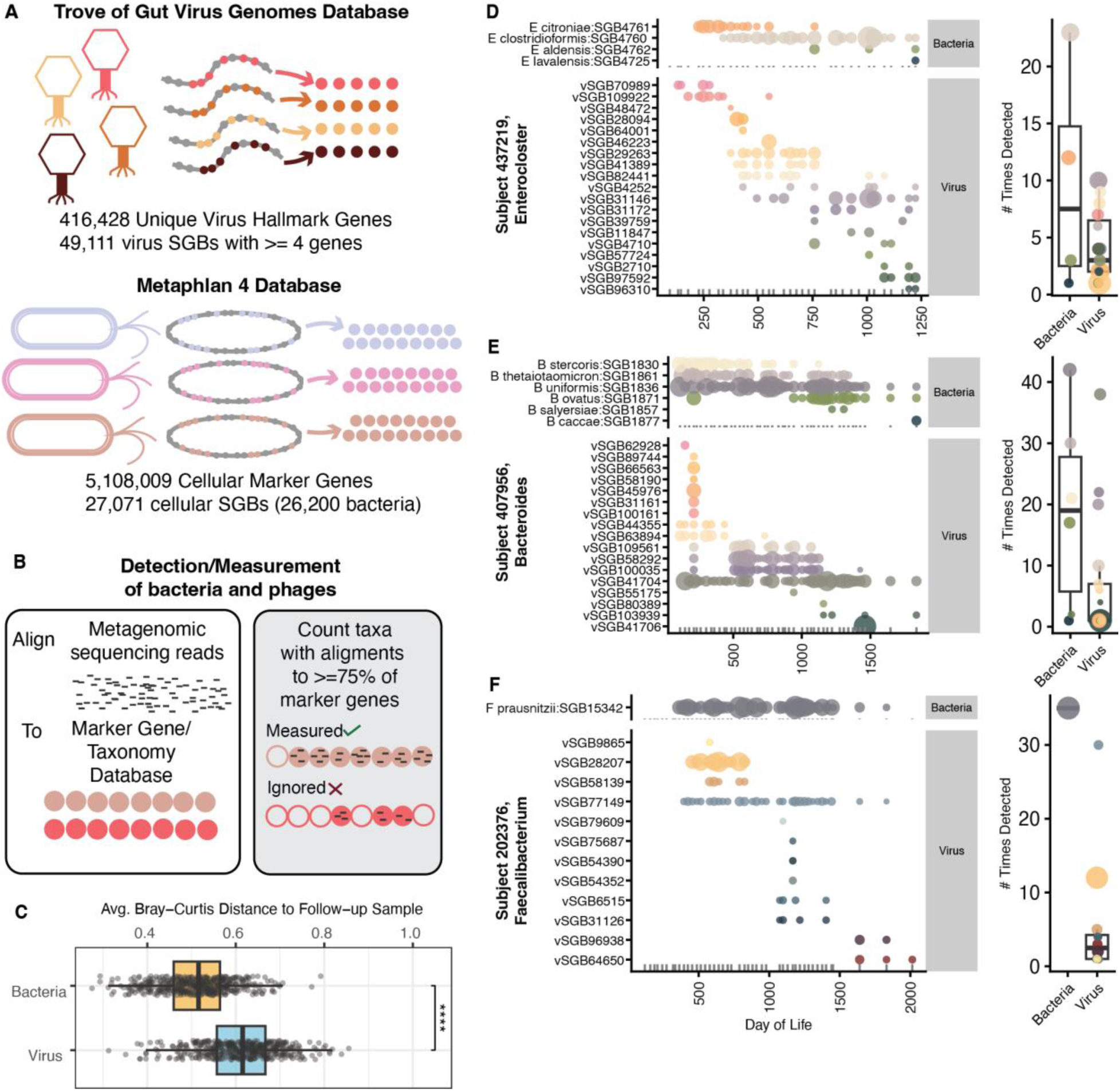
Marker-MAGu enables trans-kingdom taxonomic profiling of gut metagenomes. (A) Schematic of the Marker-MAGu database structure. (B) Marker-MAGu profiling ruleset. (C) Average Bray-Curtis dissimilarity between adjacent samples for all participants with at least 10 samples for virome and bacteriome. “****” represents p-value < 1e-04. (D - F) Examples plots of phages and their putative host bacteria prevalence as measured by Marker-MAGu.

Simulated read data from bacteria and phages show that Marker-MAGu has high specificity at all coverage levels for bacteria and viruses along with high sensitivity starting at 0.5X average read depth (Fig S2A-C). Archaea and micro-eukaryotes in the Metaphlan 4 database are detectable but only bacteria and viruses are focused on in this manuscript. Using real TEDDY WGS sequencing data, Marker-MAGu returns a consistent ratio of viral SGBs to bacterial SGBs (Fig S2D) which is not affected by the number of reads in a sample (Fig S2E). As expected, Marker-MAGu closely recapitulates Metaphlan4 abundance measurements from sequencing of a bacterial mock community, with slightly lower sensitivity and slightly better specificity (Fig S2F-G).

### Viruses Have a Higher Turnover Rate than the Bacteria in Gut Communities

To calculate viral and bacterial SGB prevalence and abundance in 12,262 WGS sequencing samples from 887 participants from the TEDDY study, sequencing reads (average of 12.0 million reads per sample) were analyzed with the Marker-MAGu marker gene approach. With WGS data, we expect to detect virus genomes inside virions, as well as dormant and actively replicating virus genomes inside host cells. Trans-kingdom taxonomic profiles for each participant show dynamic interplay between phages and their host bacteria, often revealing successive waves of phages or phage communities affecting single host bacteria (Fig 1D-F). Indeed, community change was higher from sample to sample for phage than bacteria (Fig 1C).

Across the entire dataset, 1,709 different bacterial SGBs were detected at least once, and 15,693 viral SGBs were detected at least once, with higher saturation of bacterial SGBs (Fig. 2A), with similar observations at the genus level (virus VC’s are computationally imputed genus-level clusters). While most bacterial SGBs and most viral SGBs were detected in only one or a few human participants, the skew towards rareness was much greater for viral SGBs (Fig. 2B), underpinning the individual-specific nature of the virome. Relatedly, when combining all longitudinal samples by participant, the Bray-Curtis dissimilarity of the virome was greater between participants than the bacteriome (Fig. 2C), and the alpha diversity of the virome was greater, on average, than the bacteriome (Fig. 2D).

**Figure 2.**
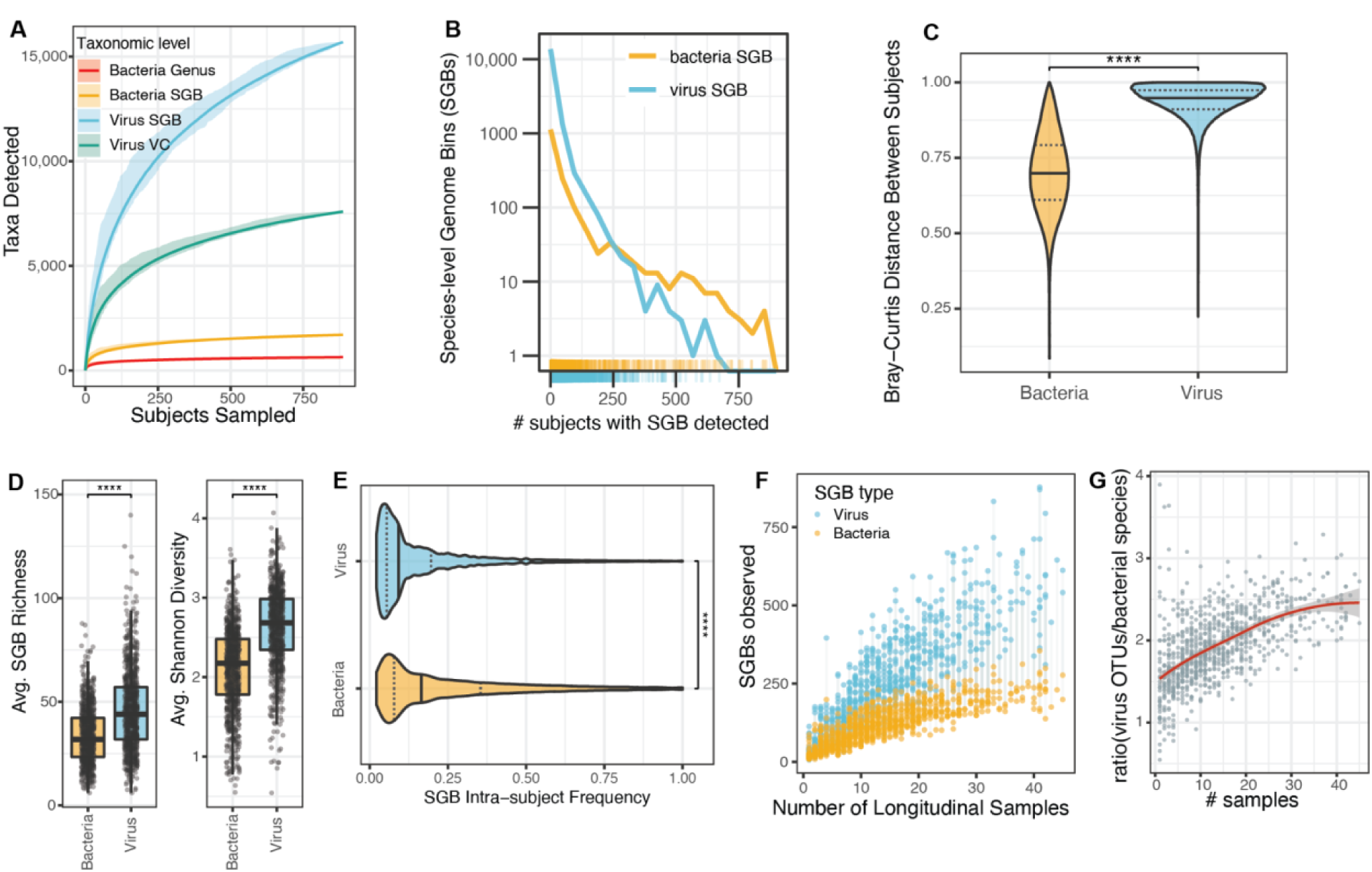
Virome and Bacteriome community change. (A) rarefaction curves of viral and bacterial taxa. (A) measurement of participant distribution of viral and bacterial SGBs. Each hash is an SGB, and the y-axis is log scale. (C) All-vs-all Bray-Curtis distance for bacteriome and virome communities after averaging microbe abundance across all available samples. (D) Alpha-diversity measurements for bacteriome and virome in gut communities. Each dot represents the average value from a participant. (E) SGB “persistence” within participants. Data was filtered to exclude participants with fewer than 10 samples. Violins represent density of values from each observation of an SGB in a participant (# times SGB detected/# total samples analyzed). Solid black line marks 50% quantile, dotted lines are 25% and 75% quantile marks respectively. (F) Cumulative number of viral and bacterial SGBs per participant. Each participant is represented by a (blue) dot for viral SGBs detected across all samples and a (gold) dot for bacterial SGBs detected across all samples connected by a gray line. (G) Ratio of cumulative viral SGBs detected vs cumulative bacterial SGBs detected. Each dot is a participant. “****” represents p-value < 1e-04.

The longitudinal nature of the data revealed that, over time, the virome has a higher participant-specific diversity and turnover rate than the bacteriome. First, it was observed that viral SGBs are detected, on average, in a smaller percentage of a participant’s samples than bacterial SGBs (Fig. 2E). Next, plotting by number of unique samples per participant, it seems new viral SGBs accumulate at a nearly linear rate, while new bacterial SGB accumulation approaches saturation (Fig. 2F). Indeed, the ratio of cumulative viral to bacterial SGBs increases from about 1.5:1 to 2.5:1 as the number of available longitudinal participant samples increases from 1 to 45.

Similar to previous reports on the bacteriome and virome in early childhood, it was observed that the alpha diversity of viruses and bacteria increased during infancy and for at least the first three years (Fig. S3A,B)^13^, and this was comparable for participants in all four countries of origin (Fig. S3C,D). Furthermore, t-SNE analysis, often used to represent relationships between high dimensional datapoints, of all samples showed samples from early infancy partitioned into many clusters while samples taken from older children converged (Fig. S3E). Samples did not partition by eventual type 1 diabetes diagnosis in this analysis (Fig. S3E). There is some debate about whether Crassvirales (a.k.a crAss-like phages) increase or decrease in abundance from infancy to early childhood^13,18^. In TEDDY, 504 putative Crassvirales genomes were detected in the Trove of Gut Virus Genomes (see Methods), and the TEDDY cohort showed a clear increase of abundance and prevalence of Crassvirales over time, with a plateau perhaps being reached 1000 days after birth (Fig. S3F).

### Common Gut Viral and Bacterial SGBs Participate in Ecological Succession in Early Childhood

To understand temporal trends in the TEDDY cohort, SGBs detected in 100 or more samples were used for further analysis. This subset consists of 446 bacterial SGBs and 1,291 viral SGBs. By calculating the relative abundance of these common SGBs from 0 - 1400 days of life across all samples, eight high-confidence “Temporal Subsets” could be separated (see Methods). Interestingly, the temporal subsets describe SGBs that peak at different host ages (Fig. 3A, B). Namely, Subsets 1 and 2 include SGBs present immediately after birth then declining thereafter. Subset 3 includes SGBs peaking at 100 - 200 days after birth. Subset 4 includes SGBs peaking at day of life 300 - 400, Subset 5 peaking at day of life 400 - 600, Subset 6 peaking at day of life 800 - 1100, and Subsets 7 and 8 continually increase over the days of life covered in this cohort. As expected based on earlier studies^11,12^, common bacteria in the earliest subsets include *Bifidobacterium breve* and *Bifidobacterium longum* whereas later subsets include *Bacteroidales* such as *Phocaeicola vulgatus*, *Bacteroides uniformis*, and *Alistipes onderdonkii* along with *Faecalibacterium species*, (Table S1).

**Figure 3.**
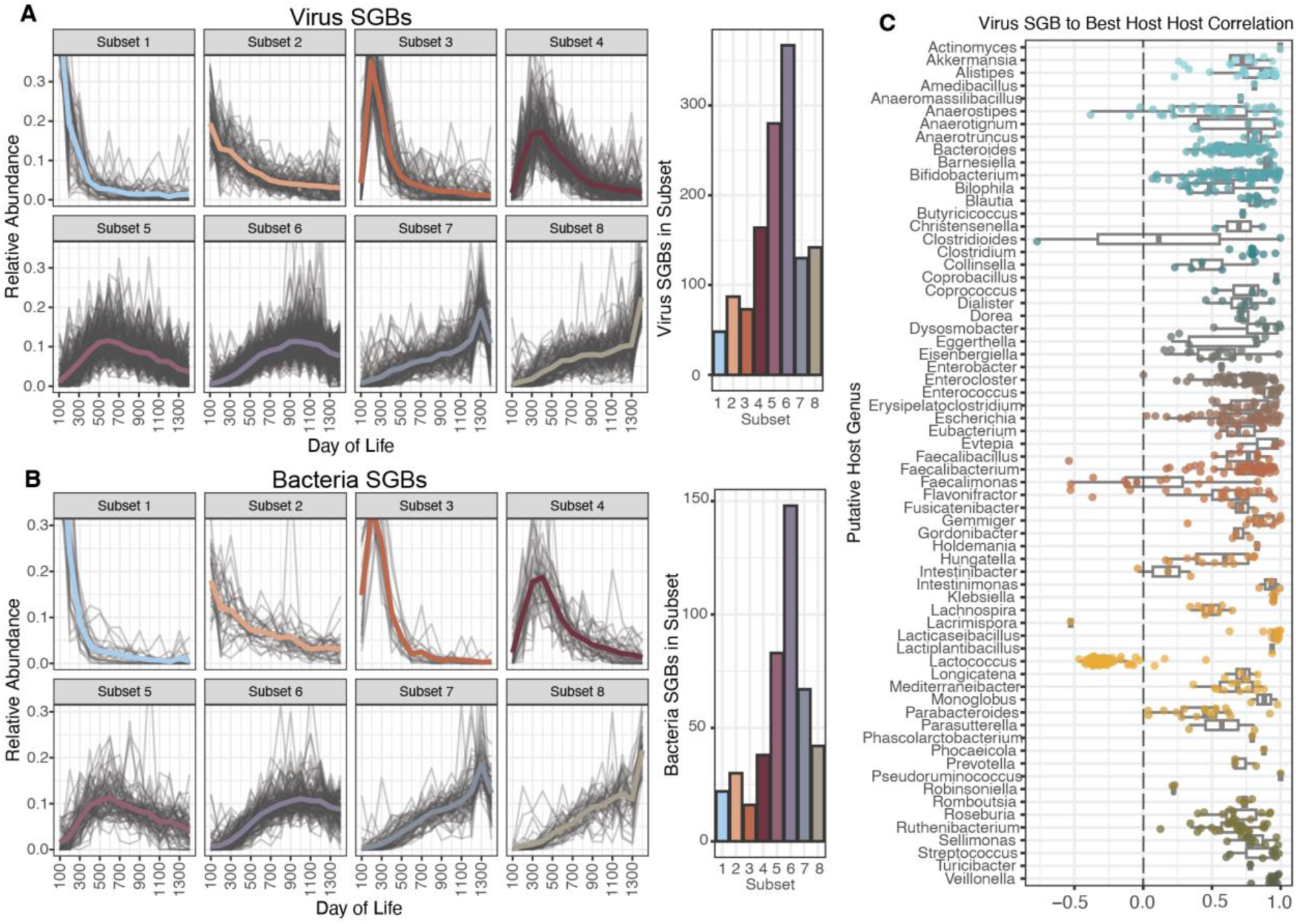
Global patterns of viral and bacterial SGBs during development. (A) Temporal Subsets of prevalent viral SGBs, drawn in separate boxes. Grey lines are individual viral SGBs and thicker colored lines are average lines. (right) Bars represent temporal cluster membership. (B) Same as (A) but with bacterial SGBs. (C) Correlation between prevalent viral and bacterial SGBs. Since host-prediction of phages is most accurate at the genus level, the temporal data from (A) and (B) was compared for each viral SGB and all bacterial SGBs from the putative host genus and the best correlation was plotted.

The viral and bacterial temporal abundance data were compared, and it was found that the viral SGBs were almost always well-correlated with their putative bacterial hosts (Fig. 3C). While this is expected of obligate parasites, the finding validates the approach used here. The major exception to this correlation was between *Lactococcus* bacteria and their phages. A definitive explanation cannot be offered, but it is notable that *Lactococcus* bacteria are commonly ingested due to their role in making dairy fermentation products such as cheese.

### Virus Abundance Data Increases Power to Discriminate Between Samples from Different Countries

The TEDDY study cohort consists of children who are vulnerable to development of type 1 diabetes (see Methods) and reside in four western countries (Germany, Finland, Sweden, and USA). To understand the how type 1 diabetes develops, each of the 114 participants in TEDDY that developed type 1 diabetes was matched with one of 114 participants who did not develop type 1 diabetes, based on geography, sex, and family history of type 1 diabetes^23^. This nested case-control study design was used here to assess microbial communities during development of type 1 diabetes. Similar to studies of bacterial community composition in the TEDDY cohort as well as metastudies of type 1 diabetes, viral and/or bacterial SGB abundance could not reliably discriminate between children who developed type 1 diabetes and those who did not (Fig. 4, right panels)^12,25^. However, random forest classifiers demonstrated that both bacteriome and virome data had discriminatory power to separate samples geographically, with virome data outperforming bacteriome data (virome better in 3/4 countries, bacteriome better in 0/4 countries), and a combination of both types of SGBs outperforming either measure alone (4/4 countries). Quantification of the most important features (SGBs) by country and day of life demonstrates geographic differences for many of these features (Fig S4).

**Figure 4.**
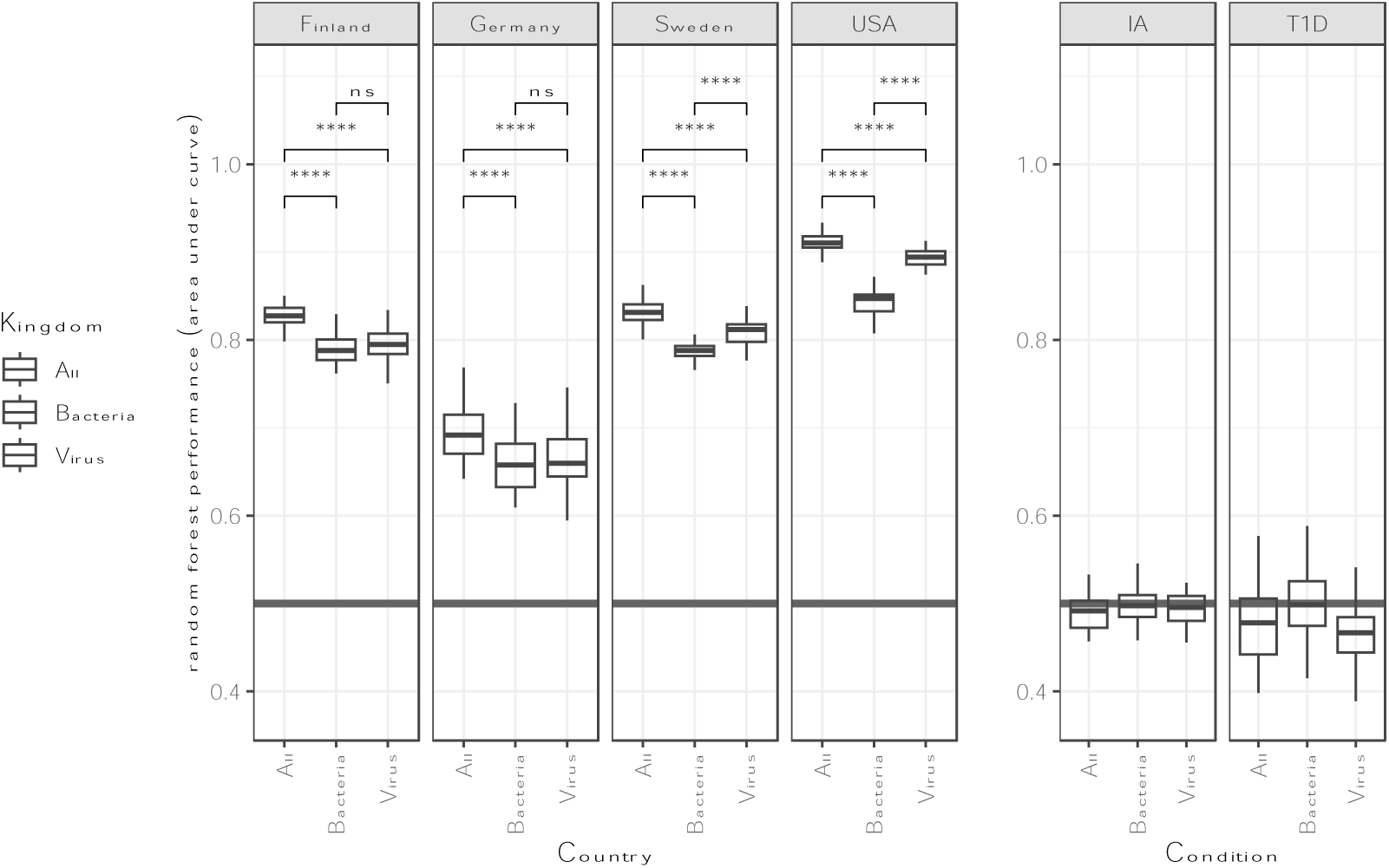
Machine Learning on Virus and Bacteria Abundance Data. Boxes represent data from 50 permutations of a random forest model on the same data split into train/test groups (70%/30%) wherein all samples from each participants were always kept in the same group. Each iteration was run with a different random seed. p-values: “ns”: 1 > p >= 0.5, “*”: 0.05 > p >= 0.01, “**”: 0.01 > p >= 0.001, “***”: 0.001 > p >= 1e-4, “****”: p > 1e-4.

### Differences in Rate of Community Change Between Groups in TEDDY Study

Analysis of viral and bacterial community change over time in each participant was conducted by calculating Bray-Curtis dissimilarity between each sample and the next, proceeding temporally. Interestingly, average Bray-Curtis dissimilarity was modestly lower in participants who went on to develop type 1 diabetes than those who did not for both virome and bacteriome measurements (Fig 5A,D). Further, when looking at comparisons across age, the trend is strongest between 400 and 700 days of life (approximately the second year of life) (Fig. 5B,E), whereas median age of diagnosis with type 1 diabetes was 2.4 years^23^. Some differences in average Bray-Curtis dissimilarity were also seen between participants from different countries (Fig 5C,F). Measuring relative abundance of the taxa from different Temporal Subsets (see Fig 3A-B), samples (particularly Day of Life 700 - 1400) from participants diagnosed with type 1 diabetes had differential abundance of Temporal Subsets 2, 5, 6, and 7 at various ages (Fig S5).

**Figure 5.**
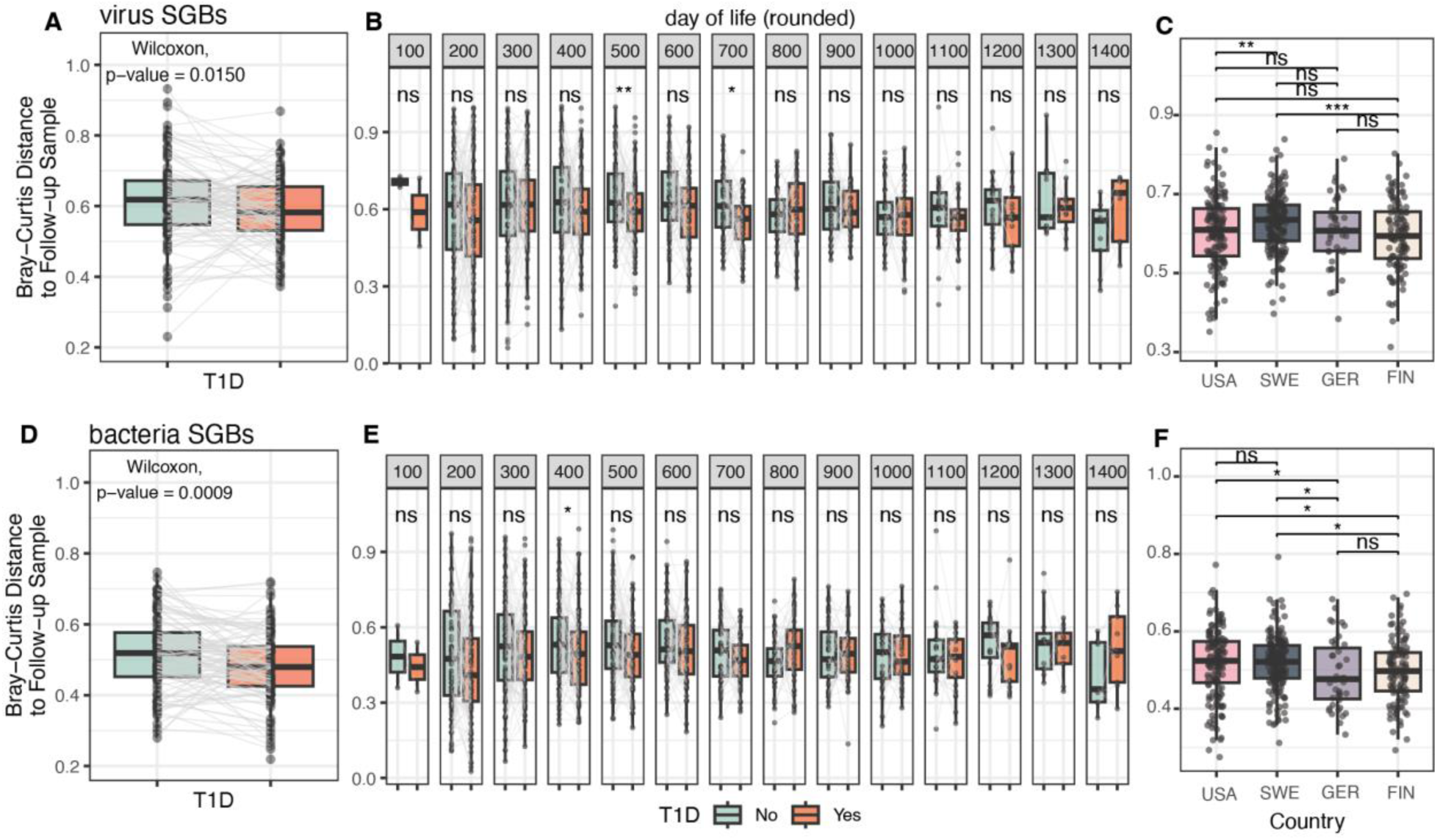
Comparison of Community Change and Diversity. (A) Virome Average Bray-Curtis dissimilarity metrics for type 1 diabetes (T1D) using nested case-control design pairing each participant who developed type 1 diabetes with a control participant. Each dot is a participant. Lines are drawn between case-control pairs. Statistical test was paired Wilcoxon test. (B) Samples are binned by day of life of collection and plotted by Bray-Curtis dissimilarity to follow up sample when both participants in pair had one or more samples from that time period. (C) Virome Average Bray-Curtis dissimilarity metrics by country (not nested case-control) (D-F) Like (A-C) but for bacteriome. (All) p-values: “ns”: 1 > p >= 0.05, “*”: 0.05 > p >= 0.01, “**”: 0.01 > p >= 0.001, “***”: 0.001 > p >= 1e-4, “****”: p > 1e-4.

## Discussion

In this study, we dissect the dynamics of the gut phages and bacteria in developing children and find individual specificity but multi-layered pattens of ecological succession. As a previous study of bacterial communities in the TEDDY cohort^12^ suggested, certain bacterial SGBs thrive at different stages of human development. This was similarly observed for viral SGBs in this study. The more complex layer of ecological succession emerged when we noticed that phages typically come and go from participants’ guts more quickly than bacteria. We then observed that many long-lived gut bacteria hosted temporally separated phages and/or phage communities during their tenure. The likely model is that newly introduced bacterial species colonize guts and succeed extant bacteria as participant diet and immune development change, (i.e. new niches emerge). With this succession, the phage communities also change, reflecting host availability. On top of this, the observation that the phages are more transitory points to an “arms race” between phages and bacteria that often results in temporary success of the phage followed by evolution of resistance in the bacterial host, rather than concurrent extinction of the phage and its host. Bacterial resistance probably emerges via point mutation, horizontal gene transfer, or introduction of a new resistant bacterial strain of the same species replacing susceptible strains. In addition, some studies suggest that strain-level replacement of bacterial species occurs frequently in the human gut^26^, and this type of switch is not detected with the abundance/detection tools used here. In a strain-level replacement, prophages would appear/disappear in the analysis while susceptibility to specific phage infections is expected to change as well.

By sampling many times longitudinally during the first years of life, it becomes clear that each participant’s gut is exposed to many more distinct phages than distinct bacteria, and this may have implications for human immune systems. Ordered arrays of antigens are highly immunogenic and likely used by B cells in detecting pathogenic viruses and bacteria^27,28^.

Phages, too, have arrayed capsid shells and tails that human immune systems may be primed to recognize and generate antibodies and memory B cells against. While some work has examined the interaction of phage and human immune systems^29^, the data presented here suggest that phages may occupy a larger share of the total antigenic surveillance space of immune systems than was previously considered. Phage antigens should be considered in future studies interested in how cross-reactivity affects inflammation, allergies, and protection against novel pathogens^30^.

While this study demonstrates that viral SGBs often quickly enter and exit the gut ecosystems of individuals, these findings do not directly contradict previous work showing that some phages can be detected in an individual’s gut over a long period of time (Fig. 1D-F)^31^.

The superiority of combining bacterial and viral SGBs over either data type alone for use in random forest models may improve the prospects for microbiome-based diagnostics. The improvement seen by including viral SGBs may be related to the geographic correlation of some bacterial strain-level phylogenies^32^. Different bacterial strains likely have unique functions/phenotypes, but also, to some extent, have distinct prophage content and phage susceptibility. However, it could also provide clues to the question “*where do our gut phages come from?*”, pointing towards acquisition dependent on a mixture of environmental conditions^11^. We also note that by keeping all the samples from a given participant together in either the testing or training group for model development likely reduced participant-specific biasing of the models (Methods).

As the attention of microbiome scientists turns towards phages, we believe it will be important to establish the rules of engagement of these viruses and their host bacteria. Phage-aware or trans-kingdom approaches, such as Marker-MAGu or recently developed Phanta^33^, will need to be adopted to this end. For example, can we identify promising phages for use in phage therapy based on how their appearance or disappearance in individuals’ guts effects potentially pathogenic bacteria. Do any phages have an effect on their host’s response to perturbations such as antibiotics, change in diet, or introgression of new bacteria? By evaluating temporal trends in the guts of developing children, we help lay the groundwork for therapeutics and diagnostics that aim to leverage the microbiome and its constituents.

## Methods

### Cohort and Study Design

In this study, whole genome shotgun data from the TEDDY study, composed of over 12,262 longitudinal samples from 887 children in 4 countries, is reanalyzed to assess phage and bacterial dynamics simultaneously. Detailed descriptions of the TEDDY Study, its cohort, and the sequencing of stool samples can be found in previous publications^12,22,23^. Briefly, the TEDDY Study is composed of six clinical research centers: three in the United States (Colorado, Georgia/Florida and Washington), and three in Europe (Finland, Germany and Sweden). The population (both cases and controls) is based on children at high risk for T1D based on their HLA genotype with 10% based on family history in addition to HLA. Collection of stool samples and associated metadata, collected using validated questionnaires, began as of 31 May 2012. Matching factors for case and control children were geographical location, sex and family history of T1D. Enrolled children were followed prospectively from three months to 15 years with stool samples collected monthly from 3 to 48 months of life, then every three months until the age of 10 years. Bacterial DNA was extracted using the PowerMag Microbiome DNA isolation kit following the manufacturer’s instructions. For whole-genome shotgun sequencing, individual libraries were constructed from each sample, pooled and loaded onto the HiSeq 2000 platform (Illumina) and sequenced using the 2 × 100 bp paired-end read protocol.^12,22,23^. The whole-genome shotgun sequencing metagenome reads are deposited in NCBI’s SRA repository under PRJNA416160. To download reads, individuals must request access to project phs001442 through the dbGaP authorization system.

### Compilation and Processing of the Trove of Gut Virus Genomes

Sequences from the Gut Virome Database, the Cenote Human Virome Database, the Metagenomic Gut Virus catalog, and the Gut Phage Database^17–20^ were downloaded and dereplicated at 95% average nucleotide identity (ANI) across 85% alignment fraction (AF) using anicalc.py and aniclust.py from the CheckV (version 0.9.0) package^34^, in line with metagenomic virus sequence community standards^24^. Exemplar sequences from each cluster/singleton from the input sequences were kept and ran through Cenote-Taker 2 (version 2.1.5)^35^ to predict virus hallmark genes within each sequence using the ‘virion’ hallmark gene database. Sequences were kept if they 1) encoded direct terminal repeats (signature of complete virus genome), one or more virus hallmark genes, and were over 1.5 kilobases or longer, or 2) encoded 2 or more virus hallmark genes and were over 12 kilobases. Sequences passing this threshold were run through CheckV to remove flanking host (bacterial) sequences and quantify the virus gene/bacteria gene ratio for each contig. Sequences with 3 or fewer virus genes and 3 or more bacterial genes after pruning/were discarded. Finally, sequences passing this threshold were dereplicated again with CheckV scripts at 95% ANI and 85% AF to yield the Trove of Gut Virus Genomes of 110,296 genomes/genome fragments each representing a viral SGB (Fig. S1).

For each sequence in the Trove of Gut Virus Genomes CheckV was used to estimate completeness, ipHOP (version 1.1.0)^36^ was used to predict bacterial/archael host genus.

Bacphlip (version 0.9.3)^37^ was run on each of the sequences predicted to be 90% or more complete to predict phage virulence.

vConTACT2 (version 0.11.3)^38^ was used to cluster viral SGBs from the Trove of Gut Virus Genomes into virus clusters. In addition to viral SGBs with vConTACT2 “Singleton” labels, viral SGBs with vConTACT2 labels “Unassigned”, “Outlier”, “Overlap”, “Clustered/Singleton” were also considered “Singletons” for downstream analysis.

### Constructing a trans-kingdom marker gene database

Hallmark genes (i.e. genes involved in replication, packaging, assembly, and virion structure) were extracted from each virus exemplar genome from the Trove of Gut Virus Genomes using HMMs from Cenote-Taker 2 (database version June 16th, 2021). Next, hallmark genes were concatenated by genome and these concatenated sequences were dereplicated at 95% ANI and 85% AF. Based on this dereplication, 70,573 virus genomes had unique virus hallmark sets based on this dereplication, indicating reduced diversity compared to whole-genome dereplication. These viruses encoded of 416,420 putative marker genes. In order to be comparable with bacterial/archaeal species-level genome bins (SGBs) from Metaphlan4^21^, which have dozens of marker genes each, detection was limited to the 49,111 viral SGBs with four or more marker genes. These genes were added to the Metaphlan4 database (version Jan21) which has marker genes from 27,071 bacteria, archaea, and micro-eukaryotes. The resulting database, which was used in these analyses, is Marker-MAGu_markerDB_v1.0.

### Marker-MAGu pipeline for marker gene-based taxonomic profiling

To quality control Illumina reads and filter out human sequences, BBDuk from BBTools was used to remove Illumina adapters, reads less than 50 nt in length and with a Q score less than 23, then read pairs were aligned to the human reference genome hg38 and phiX spike-in sequence. Unaligned read pairs were used in subsequent analyses.

Marker-MAGu is a simple pipeline. First, reads are aligned to the Marker-MAGu_markerDB_v1.0 using minimap2^39^, treated as unpaired reads. Alignments are filtered so that only reads with a single unique alignment are kept (Samtools^40^) and only reads with at least 90% identity to the reference genome across at least 50% of the read length are kept (CoverM). Read alignment information for each gene is calculated with CoverM (https://github.com/wwood/CoverM). Then, genes are grouped by taxon and, with ‘standard’ settings, only taxa with at least 75% of genes with two aligned reads are considered detected (R tyidyverse packages) (10.21105/joss.01686). Relative abundance of each taxon is then calculated. The data in this study were processed with Marker-MAGu v0.4.0 with “–detection standard”, and database v1.0.

For benchmarking, samples were processed as above and run through Metaphlan v4.0.6 with default settings.

### Identifying Crassvirales Contigs

A BLAST database was constructed from amino acid sequences of Large Terminase genes from available Crassvirales genomes in RefSeq August 17^th^, 2022. All contigs from Trove of Gut Virus Genomes were queried against this database using BLASTX with cutoffs of evalue <= 1e-5, average amino acid identity of >= 40%, and alignmnent length of >= 500 amino acids. This threshold was used as it returned 0 hits against non-Crassvirales virus genomes in GenBank.

### Diversity and Community Metrics

R package vegan was used to calculate Shannon diversity, SGB richness, and Bray-Curtis dissimilarity. Rarefaction curves were calculated using R package micropan^41^. T-SNE calculations were done using RtSNE (v0.16) (https://github.com/jkrijthe/Rtsne)

### Temporal cluster analysis

SGBs detected in 100 or more samples were analyzed to calculate average abundance from day of life 0 - 100 through day of life 1300 - 1400, in 100-day increments. Temporal clusters were calculated and assigned using R package latrend (https://github.com/philips-software/latrend").

### Random forest modeling

Random forest models were generated by using SGBs present in 1% or more samples, and by grouping all samples from each participant together either in the test or training group. Training groups were composed of samples from 70% of participants, and test groups were composed of samples from 30% of participants. A different random seed was used in each of 50 iterations of the model training/testing with python package scikitlearn^42^ to get receiver operating characteristic area under the curve (ROC AUC) and feature importance. The twelve features (SGBs) with the highest average importance over 50 iterations were chosen for supplementary plots.

### Data and reproducible script availability

The Marker-MAGu pipeline is available at https://github.com/cmmr/Marker-MAGu.

Read abundance table for bacterial and viral SGBs as well as files containing other metadata that are needed to reproduce the analyses and figures are available on <u>Zenodo</u>, doi:10.5281/zenodo.8384307.

R notebooks and Jupyter Notebooks, which can be used to reproduce each figure and analysis are publicly available on <u>github</u>.

Data was processed and parsed with R tidyverse libraries. Most figures were drawn with ggplot2 and packages ggridges (https://wilkelab.org/ggridges/), wesanderson (https://github.com/karthik/wesanderson), and nationalparkcolors (https://github.com/katiejolly/nationalparkcolors) were also used.

## Acknowledgements

The TEDDY Study is funded by U01 DK63829, U01 DK63861, U01 DK63821, U01 DK63865, U01 DK63863, U01 DK63836, U01 DK63790, UC4 DK63829, UC4 DK63861, UC4 DK63821, UC4 DK63865, UC4 DK63863, UC4 DK63836, UC4 DK95300, UC4 DK100238, UC4 DK106955, UC4 DK112243, UC4 DK117483, U01 DK124166, U01 DK128847, and Contract No. HHSN267200700014C from the National Institute of Diabetes and Digestive and Kidney Diseases (NIDDK), National Institute of Allergy and Infectious Diseases (NIAID), Eunice Kennedy Shriver National Institute of Child Health and Human Development (NICHD), National Institute of Environmental Health Sciences (NIEHS), Centers for Disease Control and Prevention (CDC), and JDRF. This work is supported in part by the NIH/NCATS Clinical and Translational Science Awards to the University of Florida (UL1 TR000064) and the University of Colorado (UL1 TR002535). The content is solely the responsibility of the authors and does not necessarily represent the official views of the National Institutes of Health.

---

**Figure S1.**
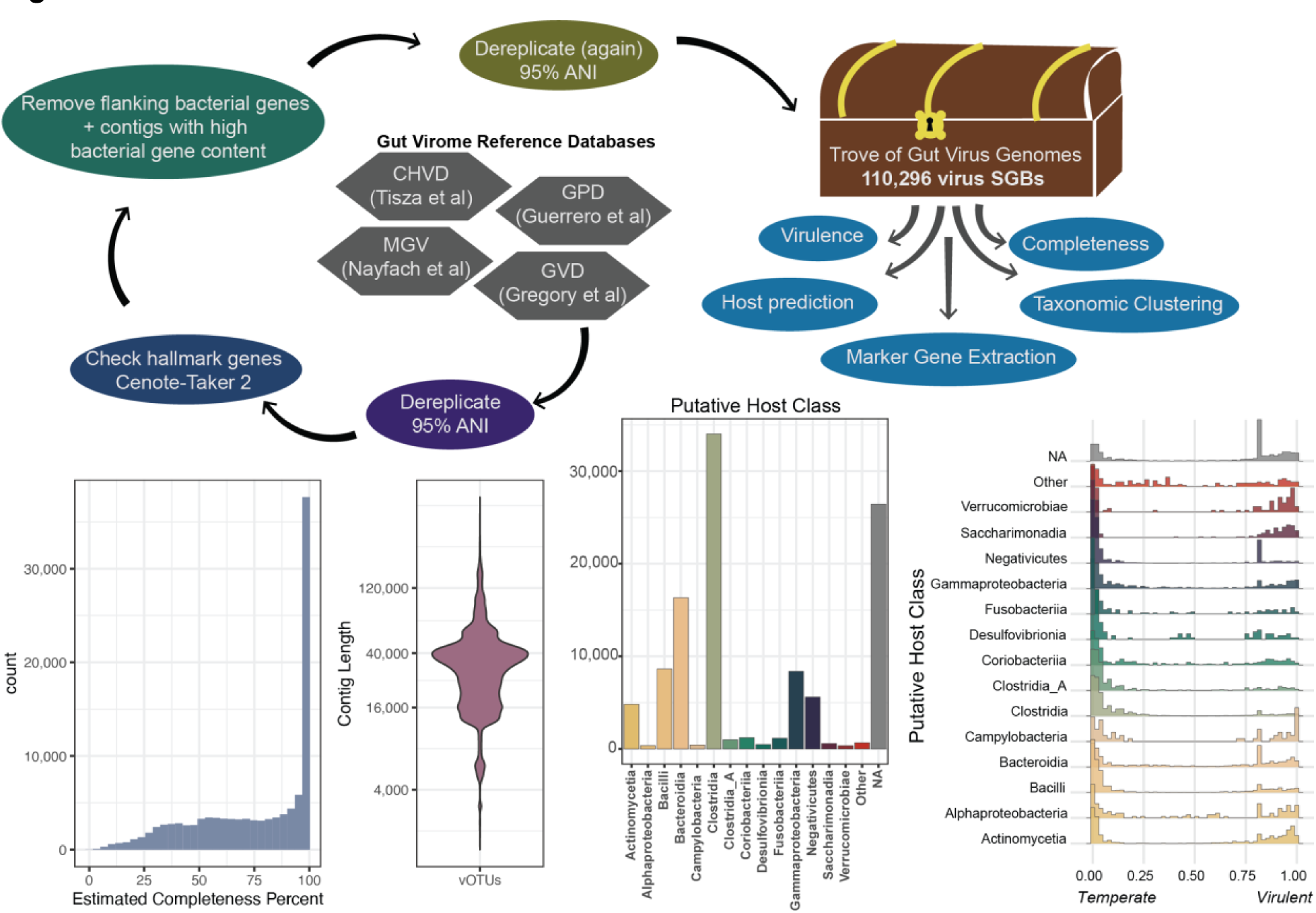
Features of the Trove of Gut Virus Genomes. (Top) pipeline to process, combine, filter and characterize extant gut virus databases. (Bottom, first from left) completeness estimate (CheckV) of 110,451 viral SGBs. (Bottom, second from left) length analysis of representative contigs for each viral SGB. (Bottom, third from left) distribution of bacterial/archael virus host prediction (drawn at class level) of viral SGBs. (Bottom, fourth from left), analysis of predicted virulence of viral SGBs for each predicted host class.

**Figure S2.**
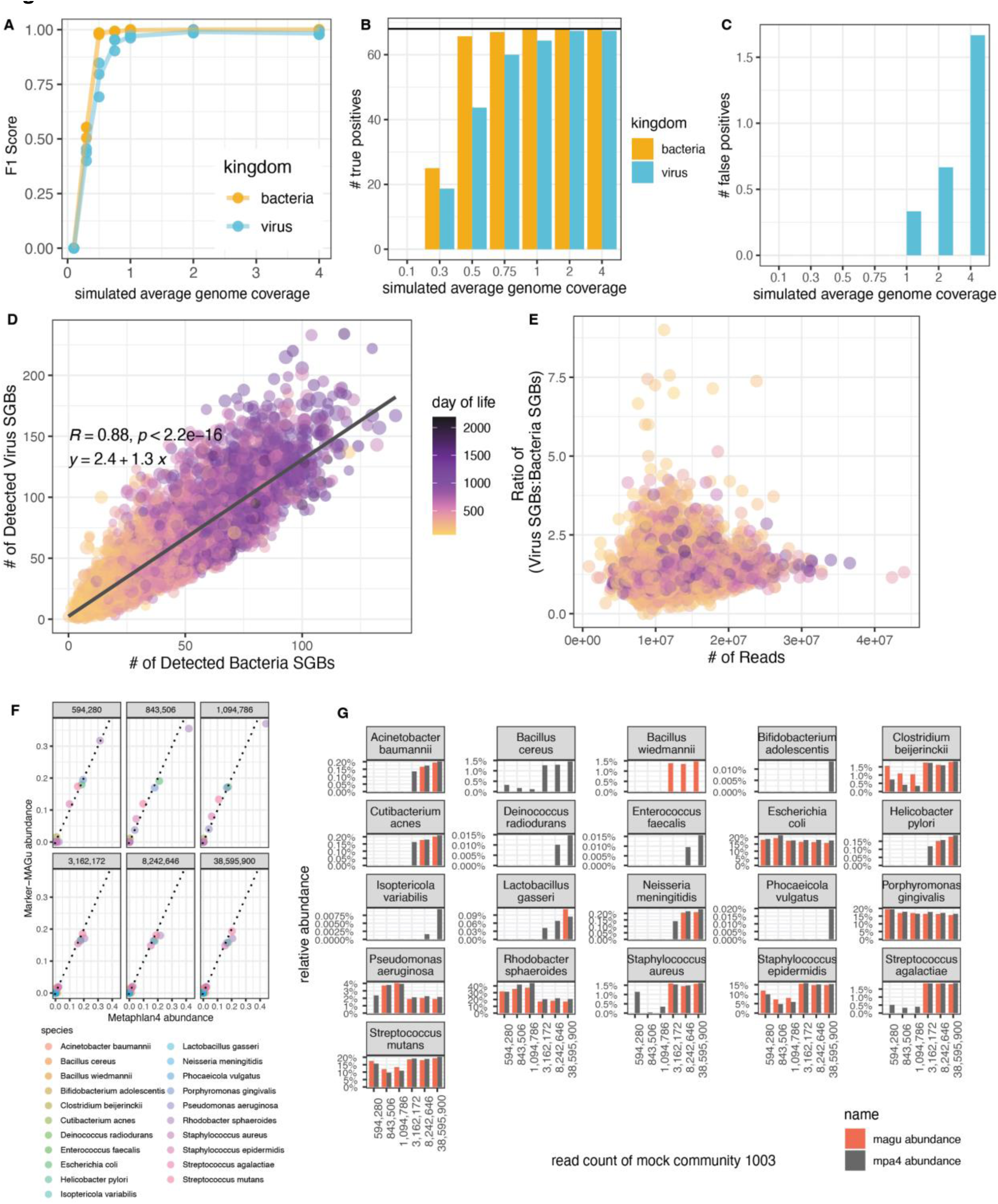
Marker-MAGu returns a consistent ratio of Viral:Bacterial SGBs across samples. (A) Simulated random reads for 68 common phage genomes and 68 common bacterial genomes were generated at 0.1X average coverage to 4X average coverage (three different random seeds each) and these reads were run through Marker-MAGu. F1 Score was calculated. (B) like but with number of true positives (68 possible). (C) like (A) but with number of false positives. (D) Scatterplot showing number of viral and bacterial SGBs detected per sample using entire TEDDY dataset. Linear relationship and pearson correlation calculated in mid-left of panel. (E) Scatterplot showing the ratio of viral SGBs:bacterial SGBs compared to number of sequencing reads for each sample, entire TEDDY dataset. (F-G) Bacterial taxa abundance measurements on sequencing of ATCC bacterial genomic DNA standard 1003 using either metaphlan4 or Marker-MAGu. Isoptericola variabilis is the only taxa not expected in the standard. *B. cereus* and *B. wiedmannii* are closely related taxa.

**Figure S3.**
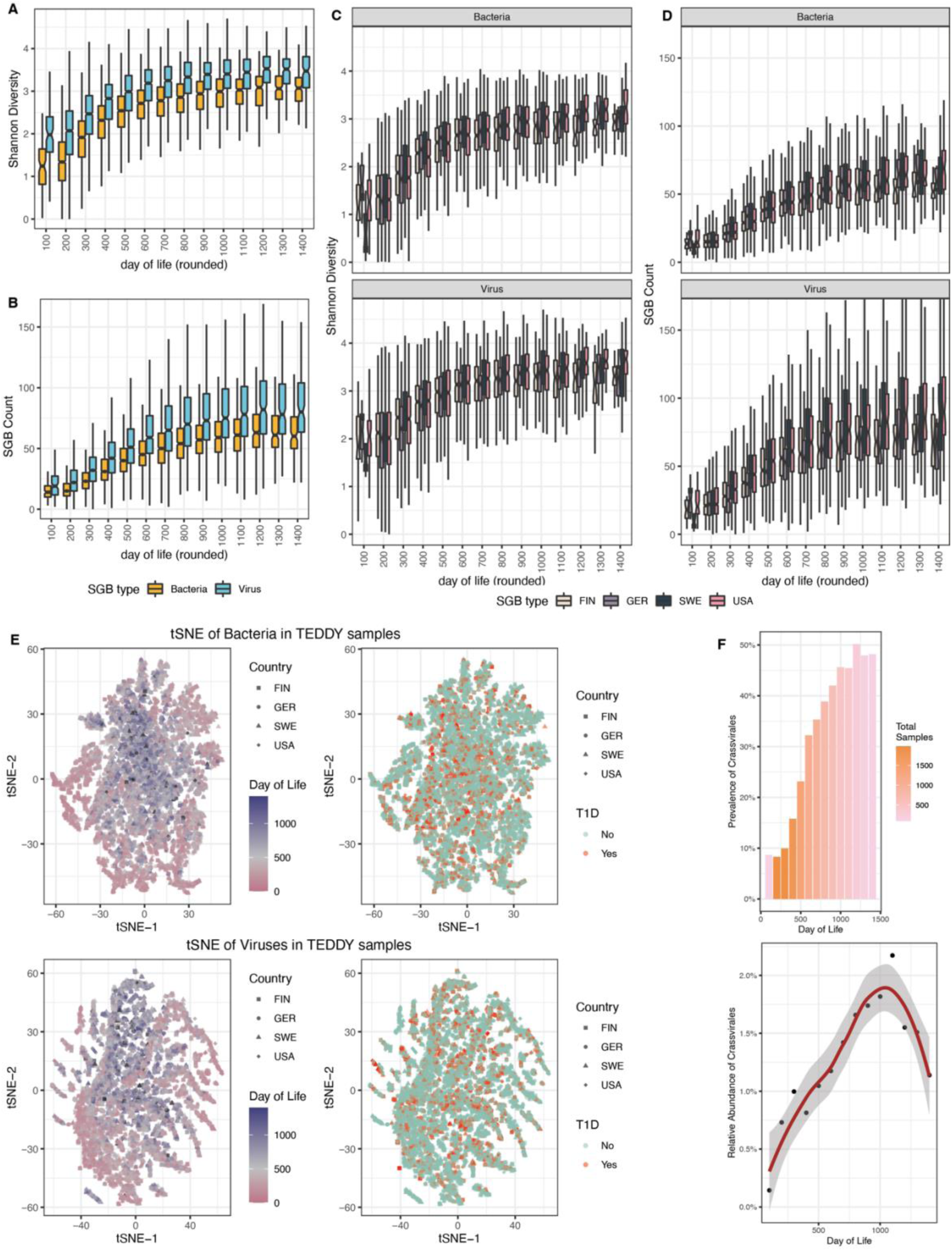
Developmental trends in the virome and bacteriome. (A) Shannon alpha diversity of samples by day of life. (B) SGB count of samples by day of life. (C) Shannon alpha diversity of samples by country and day of life. (D) SGB count of samples by country and day of life. (E) t-SNE plot of all samples bacteriome (top) and virome (bottom), colored by day of life (left) and eventual type 1 diabetes diagnosis (right). (F) Prevalence (top) and relative abundance (bottom) of all Crassvirales by day of life.

**Figure S4.**
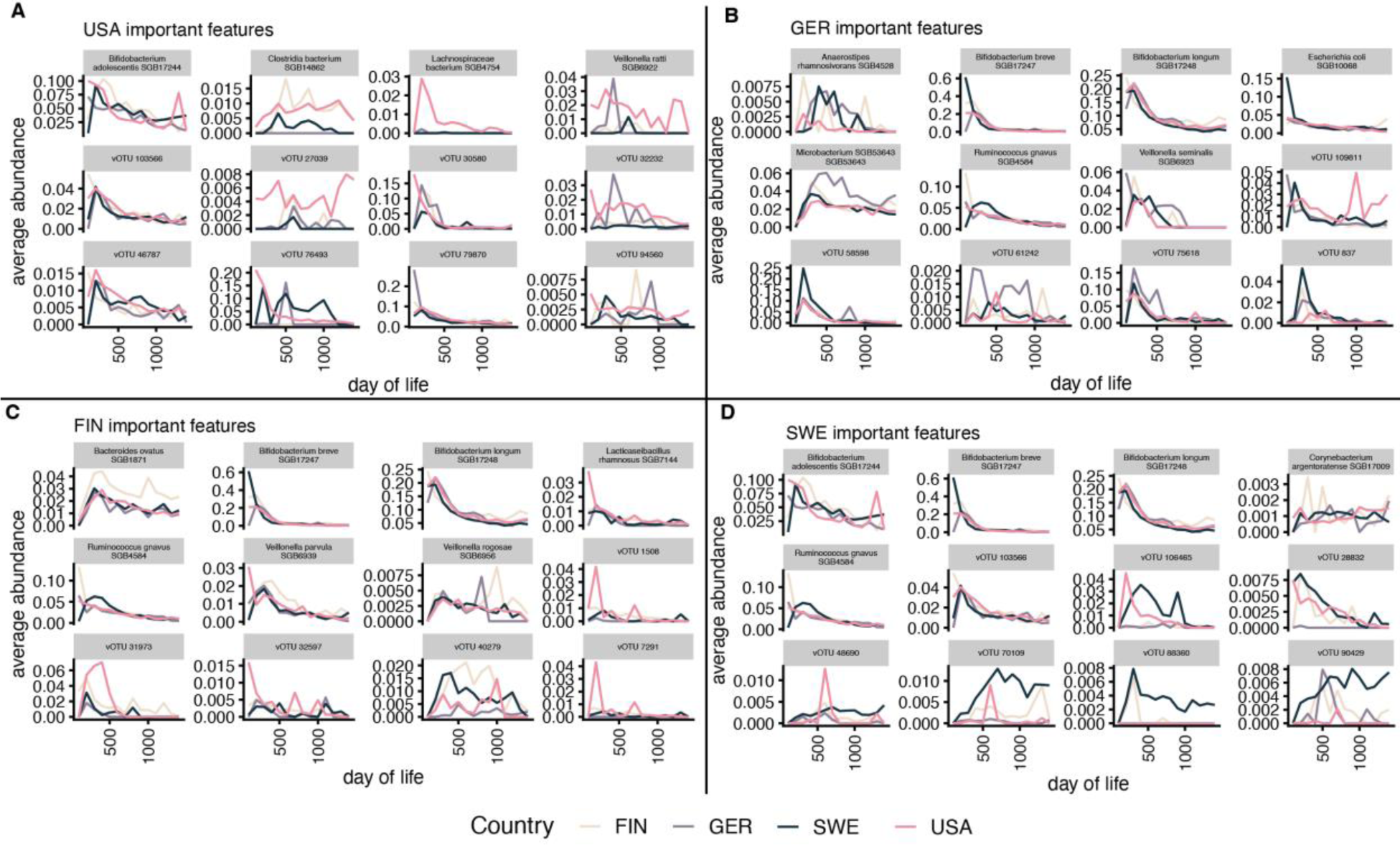
Viral and bacterial SGBs with high importance for random forest models. SGB abundance by day of life. (A) USA vs. other countries (B) Germany vs. other countries (C) Finland vs. other countries. (D) Sweden vs. other countries

**Figure S5.**
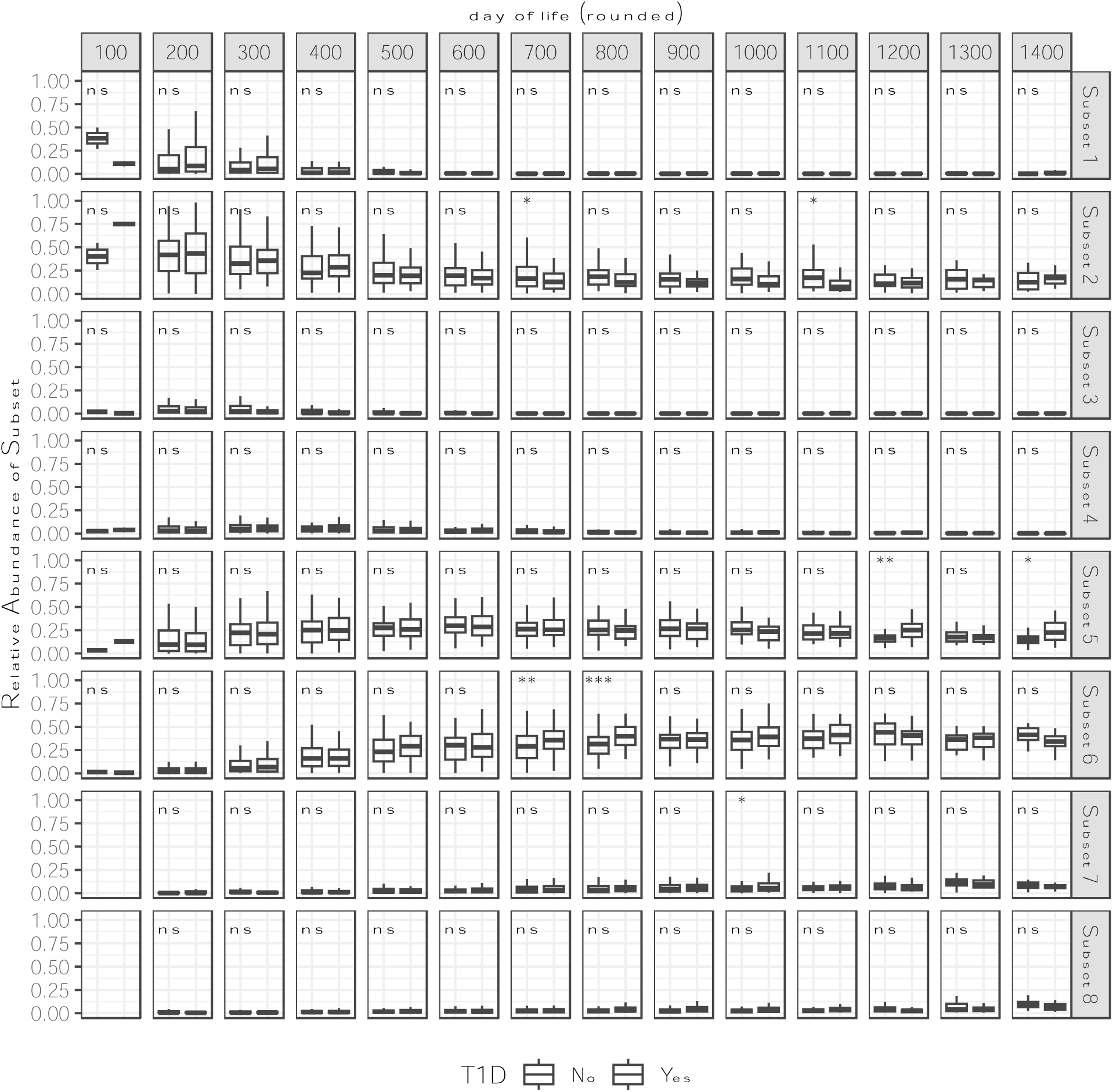
Dissimilarity for type 1 diabetes sub-groups. Abundance of prevalent Temporal Subset species (see Fig 3) for T1D and non-T1D groups across days of life. Each dot is a participant. Lines are drawn between case-control pairs. Paired Wilcoxon tests with Benjamini-Hochberg Multiple test correction. (All) p-values: “ns”: 1 > p >= 0.5, “*”: 0.05 > p >= 0.01, “**”: 0.01 > p >= 0.001, “***”: 0.001 > p >= 1e-4, “****”: p > 1e-4.

**Table S1.**
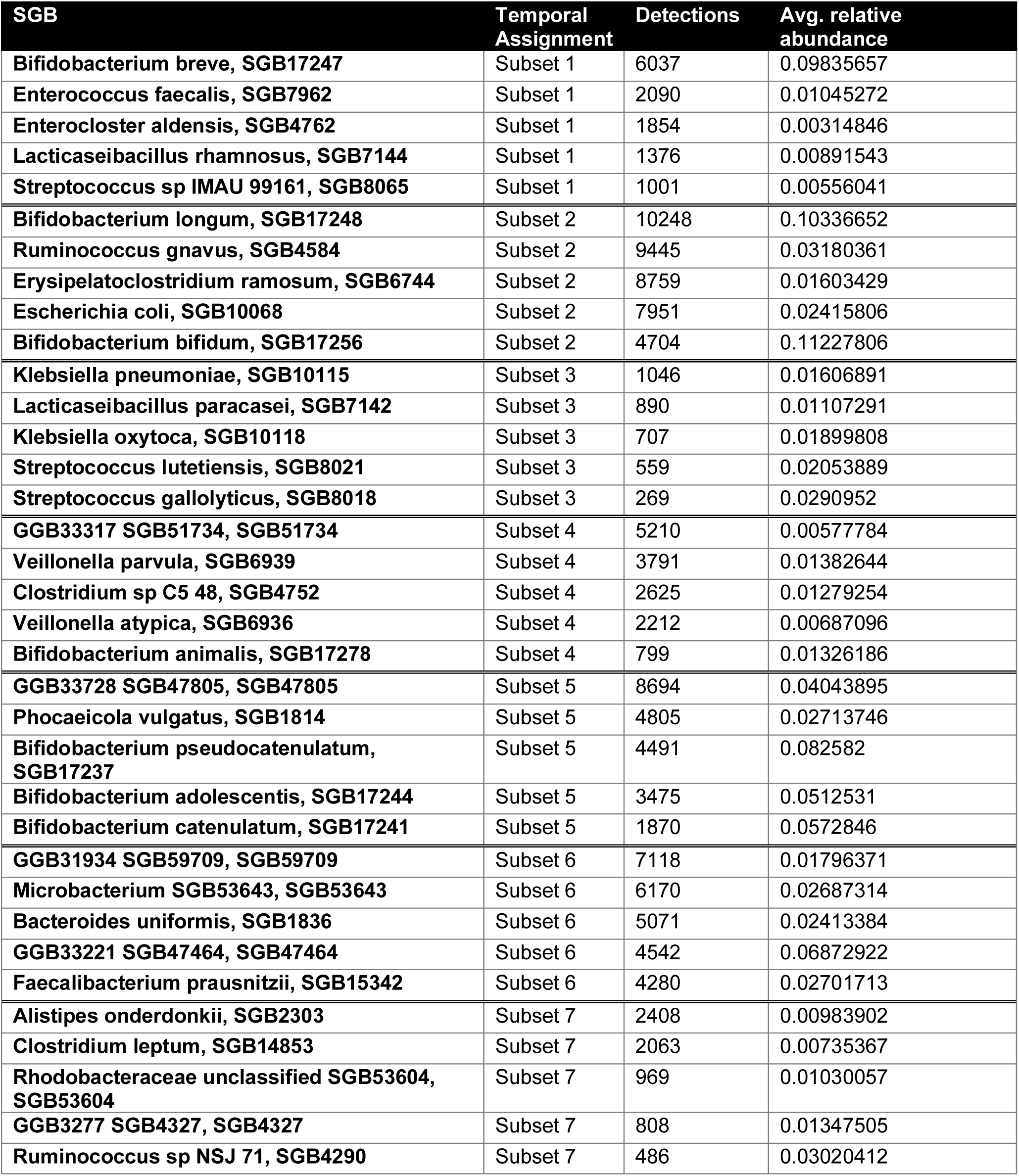
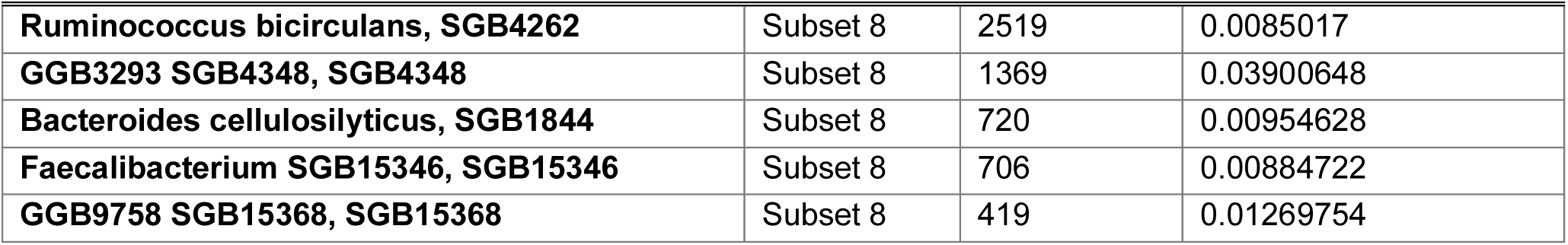
Top five most abundant bacterial SGBs in each temporal subset (product of detections and Avg. relative abundance)

